# Complex rearrangements and oncogene amplifications revealed by long-read DNA and RNA sequencing of a breast cancer cell line

**DOI:** 10.1101/174938

**Authors:** Maria Nattestad, Sara Goodwin, Karen Ng, Timour Baslan, Fritz J. Sedlazeck, Philipp Rescheneder, Tyler Garvin, Han Fang, James Gurtowski, Elizabeth Hutton, Elizabeth Tseng, Chen-Shan Chin, Timothy Beck, Yogi Sundaravadanam, Melissa Kramer, Eric Antoniou, John D. McPherson, James Hicks, W. Richard McCombie, Michael C. Schatz

## Abstract

The SK-BR-3 cell line is one of the most important models for HER2+ breast cancers, which affect one in five breast cancer patients. SK-BR-3 is known to be highly rearranged although much of the variation is in complex and repetitive regions that may be underreported. Addressing this, we sequenced SK-BR-3 using long-read single molecule sequencing from Pacific Biosciences, and develop one of the most detailed maps of structural variations (SVs) in a cancer genome available with nearly 20,000 variants present, most of which were missed by prior efforts. Surrounding the important HER2 locus, we discover a complex sequence of nested duplications and translocations, suggesting a punctuated progression. Full-length transcriptome sequencing further revealed several novel gene fusions within the nested genomic variants. Combining long-read genome and transcriptome sequencing enables an in-depth analysis of how SVs disrupt the transcriptome and sheds new light on the complexity of cancer progression.

## Introduction

Genomic instability is one of the hallmarks of cancer, leading to widespread copy-number variations, chromosomal fusions, and other sequence variations^1^. Structural variations, including insertions, deletions, duplications, inversions, or translocations at least 50bp in size, are especially important to cancer development, as they can create gene fusions, amplify oncogenes, delete tumor suppressor genes, or cause other critical changes to affect the potential of a tumor^2^. Detecting and interpreting these structural variations is therefore a crucial challenge as we try to understand the full picture of cancer genetics from cell cultures to diagnostics.

Cancer genomics has been greatly aided by the advances in DNA sequencing technologies over the last 10 years^3^. The first whole genome analysis of a cancer genome was reported in 2008^4^, and today large-scale efforts such as The Cancer Genome Atlas^5^ or the International Cancer Genome Consortium^6^ have sequenced thousands of samples using short-read sequencing to detect and analyze commonly occurring mutations, especially single nucleotide and other small variations. However, these projects have performed somewhat limited analysis of structural variations, as both the false positive rate and the false negative rate for detecting structural variants from short reads are reported to be 50% or more^7,8^. Furthermore, the variations that are detected are rarely close enough to determine whether they occur on the same molecule, limiting the analysis of how the overall chromosome structure has been altered.

Addressing this critical void, we sequenced the HER2-amplified breast cancer cell line SK-BR-3 using long-read sequencing from Pacific Biosciences. SK-BR-3 is one of the most widely studied breast cancer cell lines, with applications ranging from basic to pre-clinical research^9-11^. SK-BR-3 was chosen for this study due to its importance as a basic research model for cancer and because SK-BR-3 represents several common features of cancer including a number of gene fusions, oncogene amplifications, and extensive rearrangements. Critically, the amplifications and genome complexity observed in SK-BR-3 has been demonstrated to be representative of patient tissues as well^12^.

Taking full advantage of the benefits of long reads, we developed and applied a dual-method variant-calling approach, utilizing whole-genome assembly as well as split-read mapping to detect variants of different types and sizes. This allows us to develop a comprehensive map of structural variations in the cancer, and study for the first time how and where the rearrangements have occurred. Furthermore, combining genomic variant discovery with Iso-Seq full-length transcriptome sequencing, we discover new isoforms and characterize several novel gene fusions, including some that required the fusion of three separate chromosome regions. Finally, using the reliable mapping and coverage information from long-read sequencing, we show that we can reconstruct the progression of rearrangements resulting in the amplification of the HER2 oncogene, including a previously unrecognized inverted duplication spanning a large portion of the region. Using long-read sequencing, we document a great variety of mutations including complex variants and gene fusions far beyond what is possible with alternative approaches.

## Results

We sequenced the genome of SK-BR-3 using Pacific Biosciences (PacBio) SMRT long-read sequencing^13^ to 79.0X coverage (based on the reference genome size) with an average read-length of 9.9 kb (**Supplementary Figure 1**). In fact, we found the SK-BR-3 genome size to be much larger than the reference genome due to extensive aneuploidy. For comparison, we also sequenced the genome using short-read Illumina paired-end and mate-pair sequencing to similar amounts of coverage. To investigate the relevant performance of long and short reads for cancer genome analysis, we perform an array of comparisons in parallel using both technologies.

### Read Mapping and Copy Number Analysis

Long reads have more information to uniquely align to the genome than short reads do, resulting in overall better mapping qualities for long reads^14^ (**Supplementary Figure 2**). Using BWA-MEM^15^ to align both datasets, 69% of Illumina short paired-end reads (101bp reads, 550 bp fragment length) align with a mapping quality of 60 compared to 91.61% of reads from the PacBio long-read sequencing library (**Supplementary Figure 2, Supplementary Table 1**). We also observed a smaller GC bias in the PacBio sequencing compared to the Illumina sequence data which enables more robust copy number analysis and generally better variant detection overall (**Supplementary Figure 3**).

The average aligned read depth of the PacBio dataset across the genome is 54X, although there is a broad variance in coverage attributed to the highly aneuploid nature of the cell line (**Supplementary Figure 4,5**). Using the long read alignments, we segmented the genome into 4,083 segments of different copy number states with an average segment length of 747.0 kbp. The unamplified chromosomal regions show an average coverage of 28X, which we consider the diploid baseline for this analysis. Thus, the average copy number is approximately twice the diploid level, which is consistent with previous results characterizing SK-BR-3 as tetraploid on average^9^, and with any given locus being heterogeneous in copy number across the cell population.

Assuming a diploid baseline of 28X, the locus spanning the important HER2 (ERBB2) oncogene is one of the most amplified regions of the genome with an average of 33.6 copies (average read coverage of 470X). A few other regions show even greater copy number amplification, including the region surrounding MYC with 38 copies. Other oncogenes are also amplified, with EGFR at 7 copies and BCAS1 at 16.8 copies, while TPD52 lies in the middle of an amplification hotspot on chromosome 8 and is spread across 8 segments with an average copy number of 24.8. The locus 8q24.12 containing the SNTB1 gene is the most amplified region of the genome with 69.2 copies (969X read coverage). In addition to being the most amplified protein-coding gene in this cell line, SNTB1 is also involved in a complex gene fusion with the KLHDC2 gene on chromosome 14 **(Figure 4**).

**Figure 1.**
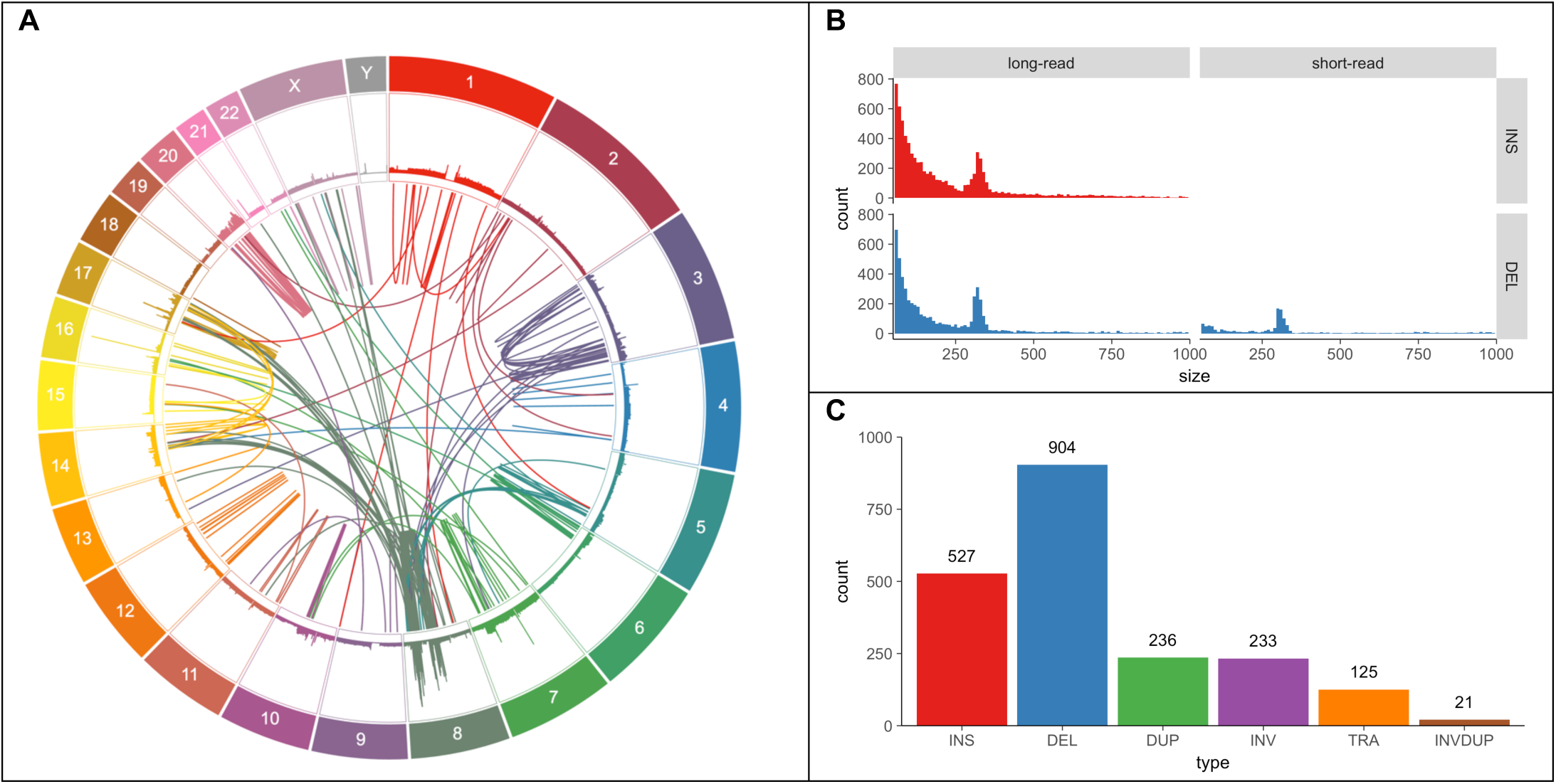
Variants found in SK-BR-3 with PacBio long-read sequencing. (A) Circos plot showing long-range (larger than 10 kbp or interchromosomal) variants found by Sniffles from split-read alignments, with read coverage shown in the outer track. (B) Variant size histogram of deletions and insertions from size 50 bp up to 1 kbp found by log-read (Sniffles) and short-read (Survivor 2-caller consensus) variant-calling, showing similar size distributions for insertions and deletions from long reads but not for short reads where insertions are entirely missing. (C) Sniffles variant counts by type for variants above 1 kbp in size, including translocations and inverted duplications.

**Figure 2.**
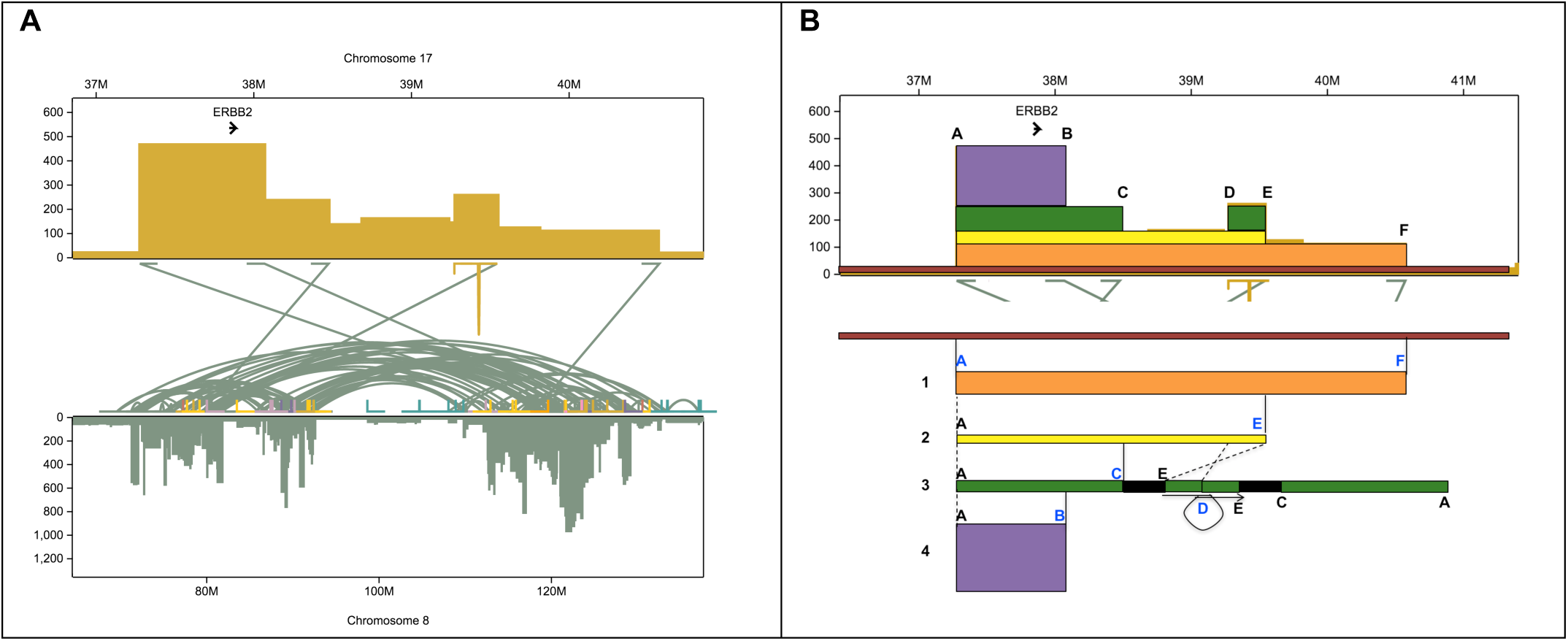
Reconstruction of the copy number amplification of the HER2 oncogene. (a) Copy number and translocations for the amplified region on chromosome 17 that includes HER2 (ERBB2). (b) Sequence of events that best explains the copy number and translocations found in this region. Segment 1 (orange) first translocated into chromosome 8, followed by the segment 2 (yellow) translocating to a different place on chromosome 8. Then the segment 3 (green) was duplicated from segment 2 by an inversion of the piece between variants D and E along with a 1.5 Mb piece of chromosome 8 that was attached at variant E, all of which then attached at variant C. The whole green segment including the 1.5 Mb of chromosome 8 then underwent an inverted duplication at variant D. The purple slice could have come from the orange, yellow, or green sequences since it only shares breakpoint A. Additionally, there is a deletion of 10,305 bp between breakpoints D and E.

**Figure 3.**
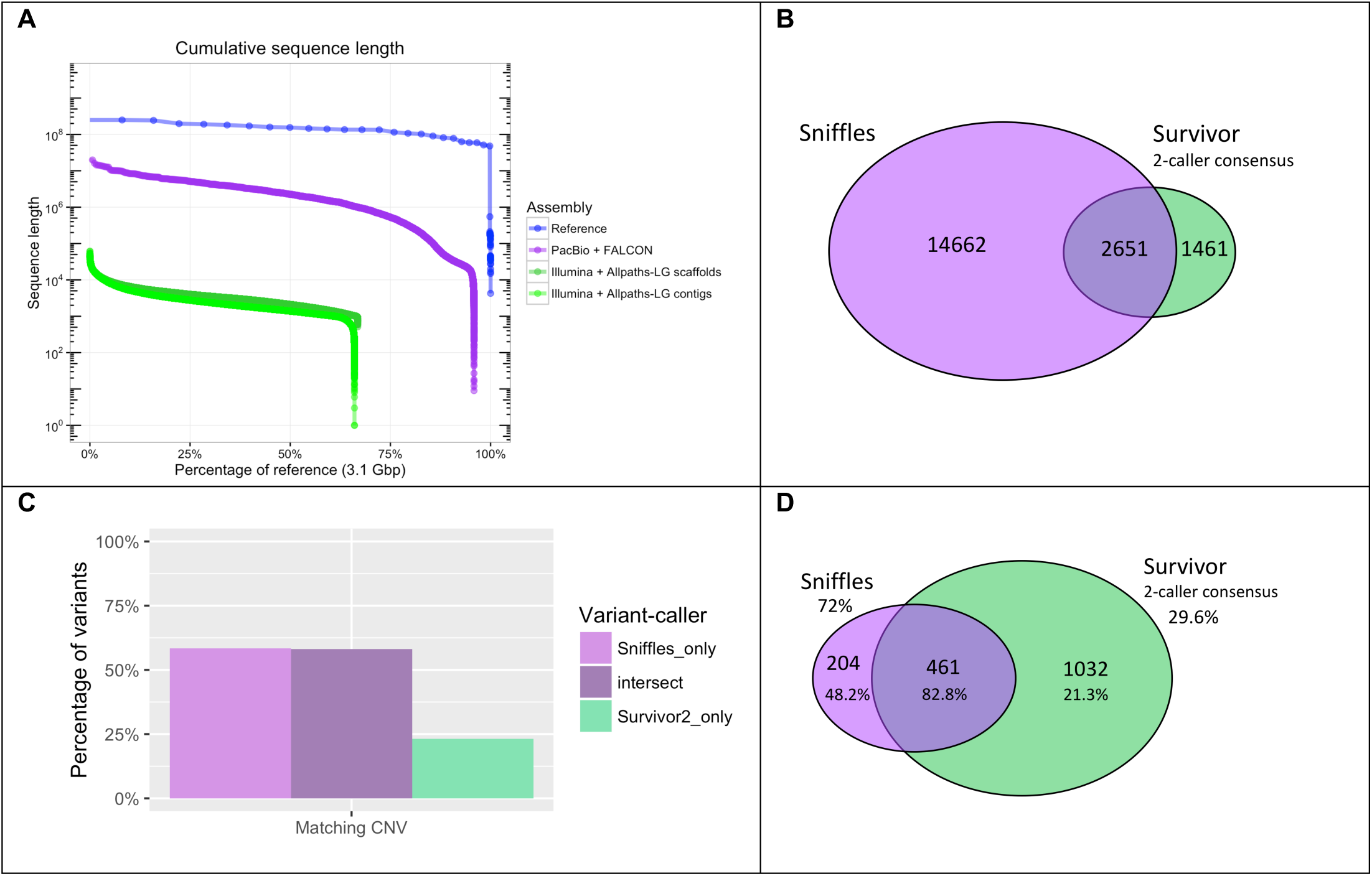
Comparing results of *de novo* genome assembly, mapping, and variant-calling between PacBio and Illumina paired-end sequencing. (A) Comparison of sequence lengths from de novo assemblies created using long reads (PacBio+FALCON, purple) and short reads (Illumina+Allpaths-LG, green), where the short-read Allpaths-LG assembly is shown as both the full scaffolded assembly (dark green) and the unscaffolded contigs (light green). The hg19 reference genome sequence lengths are shown in blue for reference. (B) Venn diagram showing the intersection of structural variants between the Sniffles call set versus the Survivor 2-caller consensus with counts indicated. (C) Percentage of variant calls in each area of Venn diagram in (B) that have matching CNV calls within 50 kbp (the smallest segment allowed in segmentation) where a CNV is a difference in copy number (long-read sequencing) between segments of at least 28X, the diploid average. (D) Venn diagram showing the intersection of long-range variants between the Sniffles call set versus the Survivor 2-caller consensus. Validation rates are shown as percentages below the counts for each category and extrapolated overall validation rates shown for Sniffles and Survivor.

**Figure 4.**
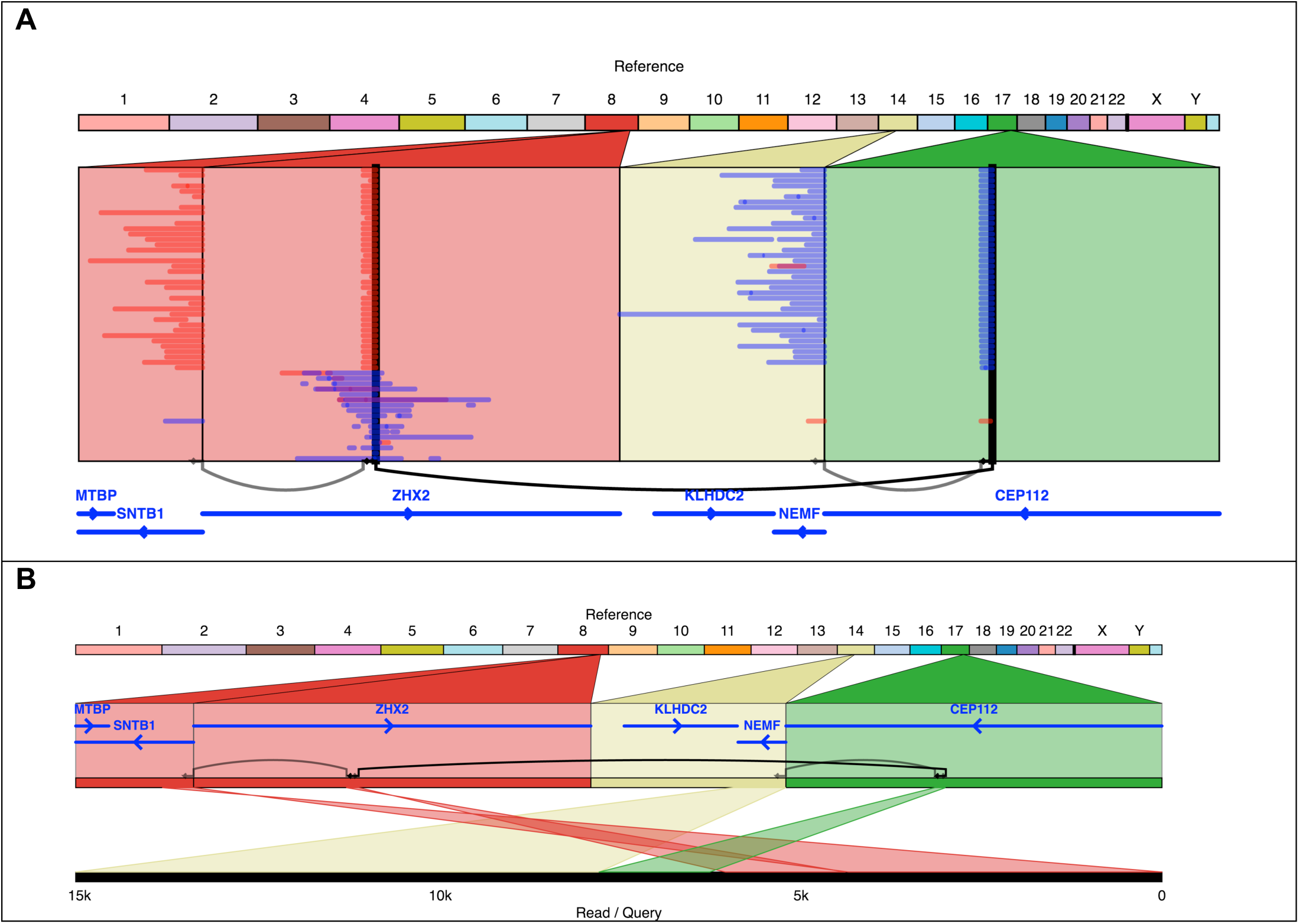
The KLHDC2-SNTB1 gene fusion in SK-BR-3 occurs through a series of three variants and is directly observed to link the two genes in several individual SMRT-seq reads (A), one of which is shown in detail in (B).

Copy number amplifications are distributed throughout the genome across all chromosomes **(Supplemental Figure 5)**. Every chromosome has at least one segment that is tetraploid or higher, and these amplified regions account for about one third (1.07 Gbp) of the genome. Extreme copy number amplifications, above 10-ploid (>140X coverage), appear on 15 different chromosomes for a total of 61.1 Mbp, with half on chromosome 8 (30.1 Mbp). There is a total of 21.3 Mbp of 20-ploid sequences across five chromosomes, with 20.0 Mbp of this on chromosome 8, and 1.3 Mbp distributed across chromosomes 17, 7, 21, and 1. In addition to containing the greatest number of base pairs of 20-ploid sequence, chromosome 8 also has 101 segments of 20-ploid sequence compared to only 4 total segments from chromosomes 7, 16, 17, and 21. Chromosome 8 thus has far higher levels of extreme copy number amplification than all other chromosomes combined.

### De novo assembly and structural variants

We generated a de novo genome assembly of SK-BR-3 from our long-read PacBio dataset using the Falcon assembler^16^. For comparison, we also assembled the genome with the widely used Allpaths-LG assembler^17^ to create a short-read assembly using a combination of an overlapping paired-end library and two mate-pair libraries. The contiguity of the long-read assembly is over one thousand-fold better than the short-read assembly, with a contig N50 of 2.4 Mbp compared to 2.1 kbp from short reads, also far surpassing the scaffold N50 of 3.2 kbp (**Figure 3A**). The high quality long-read assembly also allowed for a much more comprehensive view into structural variations compared to the short-read assembly **(Supplementary Note 1)**.

In parallel, we aligned the long reads to the reference using NGM-LR^18^ and analyzed the alignments for structural variations using Sniffles^18^ requiring at least 10 split reads to call a structural variant. Sniffles found a total of 17,313 structural variants (minimum size 50 bp), composed of 8,909 (51%) insertions, 6,947 (40%) deletions, 1,018 (6%) duplications, 279 (2%) inversions, and less than 1% total of translocations and special combined variant types **(Figure 1)**. Our work with several other genomes shows that the vast majority of these variants are correct. Including variants as small as 10 bp, there are a total of 78,776 variants are detected using the long reads, where 1,725 variants intersect 361 of the 616 genes in the COSMIC Cancer Gene census^19^. 172 of these genes are hit by structural variants (minimum 50 bp). Counting only sequences identified in Gencode as exons, a total of 58 variants intersect the exons of 46 different Cancer Gene census genes.

For comparison, we also called structural variants from a standard paired-end short-read sequencing library, using our Survivor algorithm^20^ to form a high-quality consensus call set from 3 different short-read variant callers (Manta^21^, Delly^22^, and Lumpy^23^) requiring that at least 2 of these variant-callers identified the same variant. We have found this approach reduces the false positive rate without substantially reducing sensitivity. The total number of short read structural variants in the consensus set was 4,112, composed of 2,481 (60%) deletions, 603 (15%) translocations, 580 (14%) inversions, and 448 (11%) duplications **(Figure 3b)**. Comparing the counts, the short-read consensus has a much smaller number of total variants than the long-read set (only 24%), and even when the short-read call set is expanded to include variants produced by any 1 of the callers, the total is still only 9,636 (56%) compared to the total of 17,313 for long reads. This difference is largely driven by the lack of insertions in the short-read call sets: only Manta has a limited ability to call structural insertions, with no attempts at calling these from Lumpy or Delly^21-23^. We also note that the short-read SV callers are highly enriched for false positive calls, especially false translocations (see below).

Our initial expectation was that approximately the same number of insertions and deletions would be present due to normal human genetic variation. However, the long-read variant call set has a ratio of 1.28:1 insertions to deletions. This insertional bias has been seen previously and suggests an underestimate of the lengths of low-complexity regions in the human reference genome^24^. Similar distributions of insertions and deletions, with peaks suggestive of jumping Alu elements, were found by both Sniffles and the long-read assembly-based variant-calling (**Supplementary Note 1**).

As long-range variants are of particular interest in cancer genomics, we performed several analyses specific to this subset of variants. We define long-range variants as those that are either 1) between different chromosomes, 2) connecting breakpoints at least 10 kbp apart within the same chromosome, or 3) inverted duplications. These long-range variants indicate novel adjacencies joining chromosomal regions that were originally distant in the genome. This causes novel sequence to be formed at the junction, potentially leading to gene fusions, large deletions or duplications, and other aberrant genomic features. Split reads provide a robust signal for detecting these long-range variants and chromosomal rearrangements. Within the long-read Sniffles call set, we found 665 variants in this long-range class **(Figure 1A, C)**, 125 of which were between different chromosomes. From the Survivor short-read consensus calls, 1,493 are long-range variants with 603 of these being between different chromosomes.

Focusing on the long-range variants, we analyzed the intersections between the Sniffles and Survivor (2-caller consensus) call sets. Compared to the Survivor consensus call set, Sniffles detects the same 461 and an additional 204 variants, whereas the short-read Survivor consensus detects an additional 1032 (**Figure 2B**). We selected 100 variants from each subset for PCR plus Sanger validation, with 100 calls from the intersect, 100 Sniffles calls not shared by Survivor2, and 100 Survivor2 calls not shared by Sniffles. Within each group, some variant calls could not be validated due to primer issues, so the final validation rates are calculated as successful Sanger validation counts out of the total valid attempts. As expected, the variants called by both Sniffles and Survivor had the highest validation rate of 82.8% (77/93). Of the calls unique to one method, long-read Sniffles variants have a validation rate more than twice that of the short-read variants: 48.2% (26/54) compared to 21.3% (17/80). Furthermore, extrapolating the validation rates for these subsets, the overall validation rate of Sniffles calls is 72%, while the Survivor2 calls is only 29.6%. We emphasize this is the validation rate for the most complicated long-range variants present in the genome, and our work with other long-read datasets has 94% to over 99% accuracy^18^.

Further supporting this higher validation rate of long-read variants, the Sniffles variants were also more likely to occur at the breakpoint of a copy number variant than their short-read counterparts. Specifically, 58.3% of the Sniffles unique variants show a matching copy number variant, compared to only 23.2% of the Survivor unique consensus variants, where 58.1% of the variants shared by both sets show a matching CNV (**Figure 3C**). Similar results were also found using the short reads for segmentation. The high rate of CNV matching for the shared set indicates that copy number evidence can serve as a measure of confidence in a variant call. CNV matching provides additional support for the majority of the Sniffles unique calls, though it does not exclude others that may be copy number neutral variants such as balanced translocations. The low rate of CNV support for the short-read consensus suggests that a larger proportion of these variants are either false positives or Sniffles is not sensitive enough to capture them. Reducing the threshold in Sniffles to 5 split reads (instead of the 10 split reads employed throughout this analysis) captures another 134 of the short-read consensus variants out of 1032, so there appears to be little long-read evidence of these additional variants.

### Characterization of the HER2 copy number amplification

Chromosome 8 is the most aberrant chromosome in the genome of SK-BR-3, accounting for over half of the highly amplified sequence in the genome and almost half of the long-range variants. Most of the new connections between sequences originally on chromosome 8 are clustered in three major hotspots. The HER2 (ERBB2) oncogene, originally located on chromosome 17, is amplified to approximately 32.8 copies, while most of the remainder of chromosome 17 is present in just 2 copies. The amplified region that includes HER2 contains 5 translocations (**Figure 2**) into the hotspot regions on chromosome 8, as well as an inverted duplication. Each of the 6 variants mark the site of a change in copy number, and all 6 were validated by directed PCR and Sanger sequencing. It is notable that the inverted duplication was not identified by any of the short-read variant-callers although it is clearly visible in the long-read alignments and is automatically identified by Sniffles.

The HER2 oncogene appears to have been amplified to such a great extent due to its association with the highly mutated hotspots in chromosome 8, and suggests a remarkably complex and punctuated mutational history (**Figure 2B**). The long-range variants within the amplified region containing HER2 were studied to determine whether the number of split and reference-spanning reads at each breakpoint are consistent with the copy number profile, which was found to be true for all five translocations and the inverted duplication. In order to determine the order in which these six events took place, we derived the most parsimonious reconstruction factoring in a couple of important assumptions established within population genetics. First, we assume that variants we observe have taken place only once, since it is extremely unlikely that the same long-range variant at the same two breakpoints would recur down to base-pair resolution. Second, once a variant has occurred and created an observable breakpoint, the breakpoint would not be repaired in some copies of the sequence and not others, and thus all reference-spanning reads represent an ancestral state and not a repaired breakpoint. This is an important difference from using SNPs to construct phylogenetic trees, where it is possible for a SNP to occur multiple times independently, and possibly revert back to an ancestral state. In this analysis, the long-range variants have more reliability to reconstruct the genomic history rather than SNPs because those two simplifying assumptions are extremely unlikely to be violated when two breakpoints are involved.

Given these assumptions, we can conclude that the orange segment (A-F) must have translocated first, as the other breakpoints are shared on the leftmost edge (variant A). Next the yellow segment shown in Figure 2B branched off from the orange segment because otherwise variant A must have occurred more than once. Applying the same logic, the green segment must have branched off from the yellow segment because it shares variant E, and it is not yellow branching off from green because that would violate assumption 2 by requiring that variants C and D were repaired. Variants C and D appear to co-occur in the same sequences because the copy number is the same between those two parts of the green segment and because the other sides of the variants are at breakpoints within only 1.5 Mb of each other. The only uncertainty in the ordering of events is that the purple segment could have branched off from any of the segments sharing variant A: the orange, yellow, or green segments. There is not enough information to determine which of these segments it came out of, but we can conclude that it only came out of one of them given assumption 1 that precludes multiple occurrences of the same variant.

### Complex gene fusions captured fully by long reads

In addition to genome sequencing, we performed long-read transcriptome sequencing using PacBio Iso-Seq to capture full-length transcripts. Although traditional short-read RNA-seq approaches allow isoform quantification, in many cases these reads are too short to reconstruct all isoforms, even with paired-end analysis, exon abundance, or other indirect measurements. Instead, long reads overcome such limitations by spanning multiple exon junctions and often covering complete transcripts. This makes it possible to exactly resolve complex isoforms and identify large transcripts, without the need for statistical inference^29-31^.

Iso-Seq reads were consolidated into isoforms using the SMRTAnalysis Iso-Seq pipeline^32^. In total, 1,692,379 isoforms (95.7%) mapped uniquely to the reference genome. Interestingly, the Iso-Seq RNA sequence reads indicated a total of 53 putative gene fusions each with at least five Iso-Seq reads of evidence. We further refined this candidate set using SplitThreader^33^ to exclude variants not supported by genomic structural variations. Specifically, SplitThreader searches for a path of structural variations linking the pair of genes in the putative gene fusion, requiring that the variants bring the genes together within a 1 Mbp distance. Out of 53 candidate gene fusions, SplitThreader found genomic evidence for 39 of these: 15 are the high-quality gene fusions with a genomic path between the gene bodies of at most 10 kbp shown in **Table 1**, 19 fusions overlap the first 15 (sharing the same variant and often one of the genes), and five fusions (3 non-overlapping) have paths longer than 10 kbp, leaving 14 candidate gene fusions with no genomic paths.

**Table 1.**
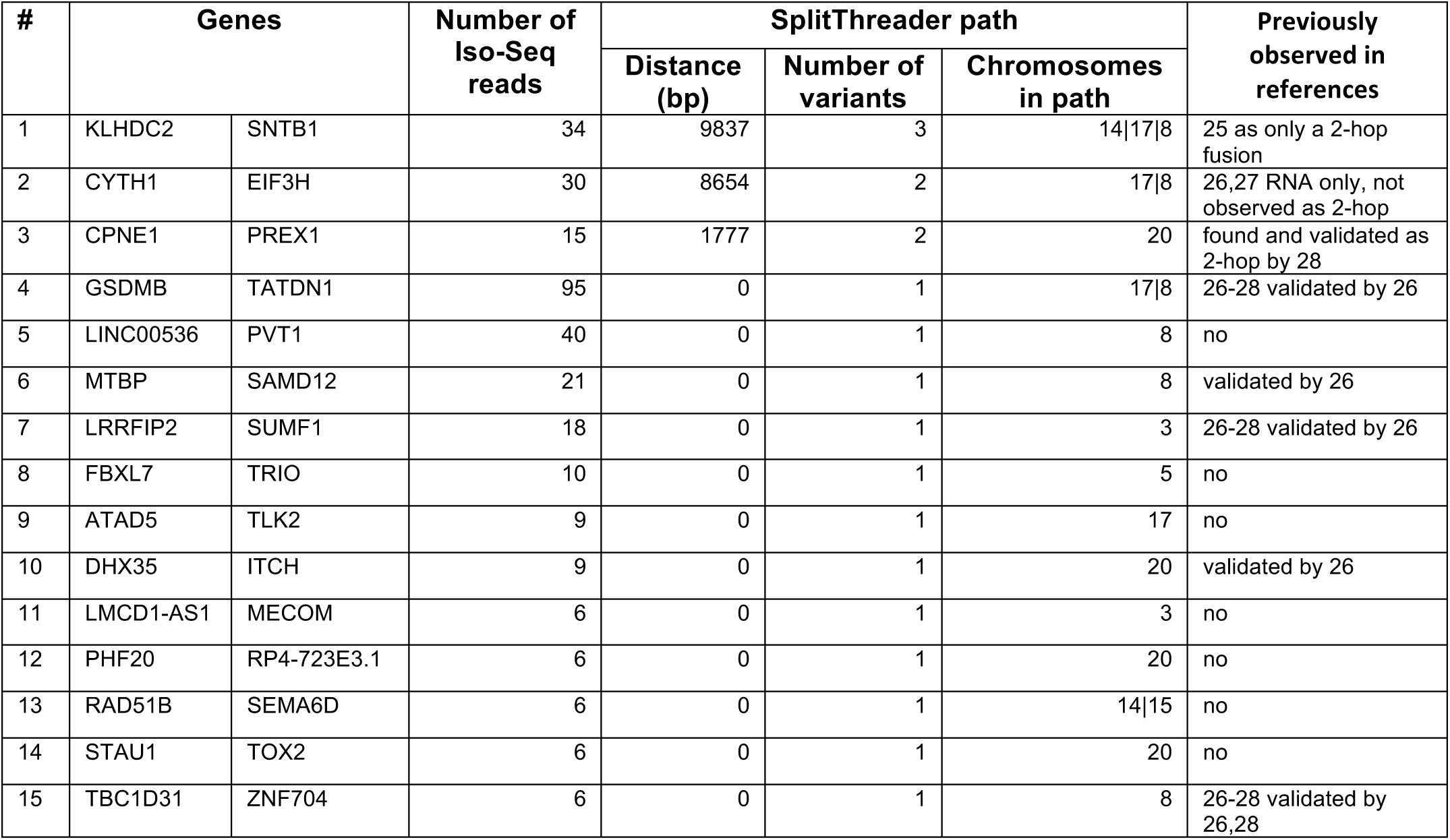
Gene fusions with RNA evidence from Iso-Seq and DNA evidence from SMRT DNA sequencing where the genomic path is found using SplitThreader from Sniffles variant calls. SplitThreader found two different paths for the RAD51B-SEMA6D gene fusion and for the LINC00536-PVT1 gene fusion. Number of Iso-Seq reads refers to full-length HQ-filtered reads. Alignments of SMRT DNA sequence reads supporting each of these gene fusions are shown in **Supplementary Note 2**.

| # | Genes |  | Number of Iso-Seq reads | SplitThreader path |  |  | Previously observed in references |
| --- | --- | --- | --- | --- | --- | --- | --- |
|  |  |  |  | Distance (bp) | Number of variants | Chromosomes in path |  |
| 1 | KLHDC2 | SNTB1 | 34 | 9837 | 3 | 14\|17\|8 | 25 as only a 2-hop fusion |
| 2 | CYTH1 | EIF3H | 30 | 8654 | 2 | 17\|8 | 26,27 RNA only, not observed as 2-hop |
| 3 | CPNE1 | PREX1 | 15 | 1777 | 2 | 20 | found and validated as 2-hop by 28 |
| 4 | GSDMB | TATDN1 | 95 | 0 | 1 | 17\|8 | 26-28 validated by 26 |
| 5 | LINC00536 | PVT1 | 40 | 0 | 1 | 8 | no |
| 6 | MTBP | SAMD12 | 21 | 0 | 1 | 8 | validated by 26 |
| 7 | LRRFIP2 | SUMF1 | 18 | 0 | 1 | 3 | 26-28 validated by 26 |
| 8 | FBXL7 | TRIO | 10 | 0 | 1 | 5 | no |
| 9 | ATAD5 | TLK2 | 9 | 0 | 1 | 17 | no |
| 10 | DHX35 | ITCH | 9 | 0 | 1 | 20 | validated by 26 |
| 11 | LMCD1-AS1 | MECOM | 6 | 0 | 1 | 3 | no |
| 12 | PHF20 | RP4-723E3.1 | 6 | 0 | 1 | 20 | no |
| 13 | RAD51B | SEMA6D | 6 | 0 | 1 | 14\|15 | no |
| 14 | STAU1 | TOX2 | 6 | 0 | 1 | 20 | no |
| 15 | TBC1D31 | ZNF704 | 6 | 0 | 1 | 8 | 26-28 validated by 26,28 |

Three of the gene fusions had no single variant directly linking the genes, but SplitThreader discovered that the genes could be linked by a series of two or even three variants. One of these, CPNE1-PREX1 had been discovered previously using RNA-seq data and validated using genomic PCR as a two-variant gene fusion^28^. We have now confirmed this by showing long reads that not only capture the two variants, but capture them together in a single read along with robust alignments to both genes (**Supplementary Figure 19**). CYTH1-EIF3H had been discovered previously with RNA-seq and been validated with RT-PCR^26^, but it was not known to be a “2-hop” gene fusion (taking place through a series of two variants) until now. This fusion was also captured in full by several individual SMRT-seq reads that contain both variants and have alignments in both genes (**Supplementary Figure 18**). Interestingly, we discovered a novel 3-hop gene fusion between KLHDC2 and SNTB1, which has been mis-reported as only taking place through two variants before^25^. We observe both the previously reported 2-hop path (600,326 bp) and this additional 3-hop path (9,837 bp), which would both result in the same gene fusion. Given the shorter distance for the 3-hop gene fusion, we were able to find direct linking evidence for the 3-hop fusion between these two genes. Strikingly, we observe 37 reads that stretch from one gene to the other through all three variants, bringing the genes within a distance of just 9,837 bp across three different chromosomes (**Figure 4, Supplementary Figure 17**). Due to the long distance between the genes through the previously reported 2-hop fusion, we believe the 3-hop fusion is more likely to produce the observed fusion transcript.

Most of the gene fusions observed are contained within a few of the most rearranged chromosomes. Four gene fusions take place within chromosome 20, which is rich in intra-chromosomal variants, while chromosome 8 is involved in six gene fusions both intra- and inter-chromosomally. The genomic variant fusing TATDN1 and GSDMB is one of the variants contributing to the amplification of the HER2 (ERBB2) oncogene. All of the gene fusions are captured fully with individual SMRT-seq reads that align to both genes. See long-read alignments spanning all 15 gene fusions in **Supplementary Note 2**.

## Discussion

Advances in long-read sequencing have produced a resurgence of reference quality genome assemblies and exposed previously hidden genomic variation in healthy human genomes^24,34,35^. Now we have applied long-read sequencing to explore the hidden variation in a cancer genome and have discovered nearly 20,000 structural variations present, most of which cannot be found using short read sequencing and many are intersecting known cancer genes. More than twice as many of the copy number amplifications could be explained through long-range variants identified by long-read sequencing compared to short-read sequencing. We further found the HER2 oncogene to be amplified through a complex series of events initiated by a large translocation into the highly rearranged hotspots of chromosome 8, where the sequence was then copied dozens of times more with further translocations and inverted duplications resolved only by the long reads. Furthermore, we find 20 additional inverted duplications throughout the genome, highlighting the importance of this underreported structural variation type. Overall, using long-read sequencing we see that far more bases in the genome are affected by structural variation compared to SNPs.

Using long-read transcriptome sequencing we capture full gene fusion isoforms, and by combining this with our genomic variant discovery, we discover several novel gene fusions in this seemingly well characterized cell line. Notably, we uncover for the first time a gene fusion that takes place through a series of three variants: KLHDC2-SNTB1 through the fusions of chromosomes 8, 14, and 17 captured fully by 37 genomic SMRT-seq reads. In a single cancer genome, we discovered three gene fusions that take place through series of two or more variants, suggesting that such multi-hop gene fusions could also be common in other cancers although they will be exceedingly difficult to discover using short-read sequencing. Conducting a similar search for multi-hop gene fusions in other highly rearranged cancers could reveal other instances of complex type of variation.

We have showed that long-read sequencing can expose complex variants with great certainty and context, suggesting that more multi-hop gene fusions, inverted duplications, and complex events may be found in other cancer genomes. Having observed complex variants such as inverted duplications with the increased informational context of long reads, the resulting variant signatures could make these events more observable even using standard short-read sequencing. However, there may be many other types of complex variations present in other cancer genomes that were not found in SK-BR-3, so it is essential to continue building a catalogue of these variant types using the best available technologies. Long-read sequencing is an invaluable resource to capture the complexity of structural variations on both the genomic and transcriptomic levels, and we anticipate widespread adoption for research and clinical practice as the costs further decline.

## Acknowledgements

We would like to thank DNAnexus for their assistance assembling the genome. This work has been supported by the NSF [DBI-1350041], the NIH [R01-HG006677], the Cold Spring Harbor Laboratory (CSHL) Cancer Center (Support Grant 5P30CA045508), the Watson School of Biological Sciences at CSHL through a training grant (5T32GM065094) from the US National Institutes of Health and by Pacific Biosciences.

## Online Methods

### Sequencing

Long-read sequencing was performed using the Pacific Biosciences Single-Molecule Real-Time (SMRT) sequencing technology with P6C4 SMRT cell chemistry. After selecting the longest subread from each polymerase PacBio read, our sequencing of SK-BR-3 yielded a mean read length of 9,872 bp, where the longest read was 71,518 bp. Total coverage of the genome is 71.9X (79.0X if redundant sequences from the same polymerase reads are included) where X refers to the number of reads that cover the average genomic base. The coverage of reads at least 10 kbp long is 51.0X, and the coverage of reads at least 20 kbp long is 13.3X. These read depth values and those in **Supplementary Figure 1B** are based on a female genome size of 3,101,804,739 bp, the total lengths of chromosomes 1-22 and X in hg19.

For short-read variant-calling, Illumina sequencing was performed on a 550bp paired-end library (2x250bp). This library produced a total of 795,942,102 reads and 64.2X genome coverage based on the same female genome size. For the short-read assembly, Illumina sequencing was performed on 180 bp paired end overlapping library (2x100 bp reads), as well as 2-3 kbp and 5-10 kbp mate-pair libraries.

### Alignment and variant-calling

The hg19 reference genome (the 1000 Genomes version) was used for all analysis. Reads were aligned to the reference using NGM-LR (v0.2.1)^18^, and Sniffles (v1.0.6)^18^ was used to call variants from long-read split alignments using the recommended parameters. Variants were called on the short-read variant-calling Illumina sequencing dataset using Manta^21^, Delly^22^, and Lumpy^23^ and a consensus was taken using Survivor^20^ with the recommended parameters, requiring two of these variant-callers to support the same variant, except where noted otherwise. Copy number segmentation was computed using SplitThreader^33^, which internally uses the DNAcopy R package for circular binary segmentation. The circos plot in **Figure 1A** was generated using Circa [http://omgenomics.com/circa]. Cancer gene intersects were determined using bedtools pairtobed to intersect Sniffles variants down to 10bp in size with the GENCODE hg19 annotation^36^ and filtered by matches to the COSMIC Cancer Gene census^19^.

### Mapping comparison

In order to compare the mappability of long and short reads, we aligned both the paired-end Illumina sequencing and the PacBio long-read sequencing datasets to the hg19 reference genome using BWA-MEM^15^. The Illumina sequencing was performed using a 550bp paired-end library with each read being approximately 250bp of sequence. We trimmed these reads to 101bp and compared both of these against the PacBio dataset. All three read sets were aligned using default parameters, except that the PacBio reads were aligned using the pacbio alignment mode in BWA-MEM (-x pacbio). The maximum mapping quality in BWA-MEM is 60, and the minimum is 0. Using the same aligner allows us to better compare mapping quality scores for the reads. We analyzed the mapping quality from each type of sequencing in two different ways, by individual reads and by binned windows in the genome. First, we selected the best alignment by mapping quality for each read and counted the number of reads in each category: mapping quality of 60, mapping quality between 1 and 59, mapping quality of 0, or unmapped. Alignment of PacBio sequence reads resulted in 91.6% of reads mapping with a mapping quality of 60, compared to only 71.2% of Illumina reads (69.0% of the 101bp trimmed reads). A greater fraction of reads from PacBio long-read sequencing map uniquely to the genome compared to short reads from Illumina sequencing (**Supplementary Figure 2, Supplementary Table 1**).

In order to determine the effect of GC content (the fraction of guanine and cytosine as opposed to adenine and thymine in a particular region), we counted the GC-fraction of each 10 kbp window in the genome, excluding those containing Ns in the reference, and calculated the read coverage from each dataset. The read depth of each 10kb bin is shown in **Supplementary Figure 3** on a log scale versus the GC fraction, along with a Lowess fit for each dataset. There is a higher GC-bias in the Illumina datasets compared to the PacBio data set, as seen by a lower read depth in bins with a higher GC fraction, while for SMRT sequencing there is a much lower bias.

To determine the read depth per chromosome, we used bedtools to find the distribution of read depth for each chromosome for the PacBio, Illumina 250bp, and Illumina 101bp datasets. These are shown as a violin plot of Gaussian kernel distributions for each chromosome in **Supplementary Figure 4**. The shapes of the distributions are largely consistent between sequencing technologies.

### Assembly

The assembly was generated from the SMRT-sequencing reads using FALCON^16^ on the DNAnexus platform. To produce a short-read assembly for comparison, the overlapping fragment library and the mate libraries Illumina reads were assembled using Allpaths-LG^17^. For assembly-based variant-calling in **Supplementary Note 1**, alignment of the PacBio assembly contigs and Illumina assembly contigs (not scaffolds) to hg19 was computed using MUMmer^37^, and Assemblytics^38^ was used to call variants.

### Iso-Seq and gene fusion analysis

PacBio Iso-Seq sequencing was performed in four size batches (0.8-2kb, 2-3kb, 3-5kb, and 5-10kb). The Iso-Seq data were processed using the SMRTAnalysis (version 2.3) Iso-Seq pipeline, which generated 441,932 high-quality (HQ), full-length Quivered consensus sequences, which were then aligned using GMAP^39,40^ to hg19. The GMAP alignments were filtered using quality scores from BWA-MEM^15^ alignments by removing any reads that in BWA-MEM have alignments below a mapping quality of 60. The remaining GMAP alignments were used for gene fusion detection using TOFU. Aligned fusion transcripts identified by TOFU were intersected with the GENCODE hg19 annotation^36^, and the total number of full-length reads supporting fusions between each pair of genes was counted. All putative gene fusions with at least 5 full-length Iso-Seq reads from TOFU were input into SplitThreader^33^ to identify those with any combination of long-range variants that place the genes within 100 kbp of each other. Gene fusion alignments were visualized and figures generated using Ribbon^41^.

## Data Availability

The raw reads, alignments, assemblies, and supplemental code are available at https://github.com/schatzlab/SKBR3

